# iPSC-based modeling of THD recapitulates disease phenotypes and reveals neuronal malformation

**DOI:** 10.1101/2022.02.24.481741

**Authors:** Alba Tristán-Noguero, Irene Fernández-Carasa, Carles Calatayud, Cristina Bermejo-Casadesús, Leticia Campa, Francesc Artigas, Rosario Domingo-Jiménez, Salvador Ibáñez, Mercè Pineda, Rafael Artuch, Ángel Raya, Àngels García-Cazorla, Antonella Consiglio

## Abstract

Tyrosine hydroxylase deficiency (THD) is a rare genetic disorder leading to dopaminergic depletion and early-onset parkinsonism. Affected children present with either a severe form that does not respond to L-Dopa treatment (THD-B), or a milder L-Dopa responsive form (THD-A). We generated induced pluripotent stem cells (iPSCs) from THD patients that were differentiated into dopaminergic neurons (DAn) and compared with control-DAn from healthy individuals and gene-corrected isogenic controls. Consistent with patients, THD iPSC-DAn displayed lower levels of DA metabolites and reduced TH expression, when compared to controls. Moreover, THD iPSC-DAn showed abnormal morphology, including reduced total neurite length and either an abnormal TH proximodistal gradient (THDA), or neurite arborization defects (THDB). Treatment of THD-iPSC-DAn with L-Dopa rescued the neuronal defects and disease phenotype only in THDA-DAn. Interestingly, L-Dopa treatment at the stage of neuronal precursors could prevent the alterations in THDB-iPSC-DAn, thus suggesting the existence of a critical developmental window in THD. Our iPSC-based model recapitulates THD disease phenotypes and response to treatment, representing a promising tool for investigating pathogenic mechanisms, drug screening, and personalized management.

## Introduction

Tyrosine hydroxylase deficiency (THD, OMIM 191290) is an ultra-rare autosomal recessive disorder caused by mutations in the gene that codes for the tyrosine hydroxylase (TH) enzyme, which catalyzes the rate-limiting step in the biosynthesis of dopamine, leading to dopamine brain depletion. Approximately 80 cases of THD have been described worldwide (Willemsen *et al,* 2010; Dong *et al,* 2020), but it is believed that misdiagnosis rates are not negligible.

More than 40 disease-causing mutations in the human TH gene have been described (Willemsen *et al,* 2010; Furukawa & Kish, 2017), among which the most frequent ones, are p.Arg233His and p.Leu236Pro (Willemsen *et al,* 2010). Defects in TH gene can cause intellectual disability, tremor, chorea, oculogyric crisis, eyelid ptosis, diurnal fluctuation of signs and autonomic dysfunction. However, patients can present with a wide phenotypic spectrum without clear correlation between genotype and phenotype, as the same mutation, occurring in homozygosis or compound heterozygosis, can be associated with mild or severe phenotype (Willemsen *et al,* 2010; Fossbakk *et al,* 2014). Children affected by the mild form of this disorder (Type A) typically develop a progressive hypokinetic-rigid syndrome and dystonia with an onset in infancy or childhood, while the severe form of TH deficiency (Type B) cause an early onset severe encephalopathy with global developmental delay and severe parkinsonism (Willemsen *et al,* 2010; Furukawa & Kish, 2017).

Since THD leads to dopamine deficiency in the CNS, treatment with L-dopa and Carbidopa represent the strategy of first choice. Type A THD patients usually respond to this treatment achieving normal cognitive and motor milestones, although cognitive development may be moderately abnormal in some patients, despite appropriate treatment (Willemsen *et al,* 2010). Regarding the THD severe form, most of these Type B patients do not show improvement of encephalopathy or motor abilities when treated with levodopa (Willemsen *et al,* 2010; Furukawa & Kish, 2017). Moreover L-Dopa supplementation in those patients is often associated with secondary effects, such as levodopa-induced dyskinesias (Pons *et al,* 2013).

Therefore, early diagnosis and the identification of reliable biomarkers of THD as well as the establishment of new therapies, represents an urgent medical need. However, the understanding of the molecular basis of THD and the heterogeneity within and between different forms of the disease is still poor due to the low prevalence of the disease and the few studies with patients’ samples which have provided only limited information (Dong *et al,* 2020).

Murine models of THD have been developed and represent valuable models to evaluate the safety and efficacy of therapies; nevertheless, they fail to fully recapitulate the spectrum of pathological manifestations observed in patients (Kobayashi et al, 1995; Althini et al, 2003; Tokuoka et al, 2011; Rose et al, 2015; Korner et al, 2015; Zhou et al, 1995). The THD mouse model (Korner *et al,* 2015) carrying Th p.Arg203His mutation (equivalent to p. Arg233His in humans, the most recurrent mutation producing THD), show alterations, such as a decrease in brain dopamine and low brain TH. However, these mice do not reflect the patient heterogeneity in motor dysfunction, biochemical phenotype and variability in response to L-DOPA treatment (Willemsen *et al,* 2010; Fossbakk *et al,* 2014), which would imply that THD is a heterogeneous autosomal recessive disease, in which different steps of the catecholaminergic systems can modulate the human phenotype.

Currently used THD cellular model include rat PC12 cells where different TH human mutations have been introduced, that do not resemble relevant disease cell type or patient metabolic and functional features (Hole *et al,* 2015). In a recent study, using a post-mortem brain tissue of a Type B patient (TH p.Arg328Trp and p.Thr399Met), low expression levels of dopaminergic proteins including TH, AADC, VMAT1, VMAT2, D1DR and D2DR and an altered expression of GABAergic and glutamatergic proteins, have been described (Tristán-Noguero *et al,* 2016). However, the effect of TH mutations in dopaminergic neurons (DAn) or the major pathogenic mechanism underlying the different phenotypes among THD type A and type B, remain unclear. Thus, there is a strong need for human-derived experimental models to explore THD pathogenesis and test new therapeutic strategies.

Here, we reprogrammed fibroblasts from THD patients with mild (Type A; TH p.Arg233His) and severe (Type B; TH p.Arg328Trp and p.Thr399Met) forms of the disease (hereafter referred to as THDA and THDB, respectively). Induced pluripotent stem cells (iPSCs) were differentiated into patient-specific DAn, which were compared in terms of morphology, dopamine metabolism, and treatment responsiveness with DAn from healthy individuals and gene-corrected isogenic controls. THD iPSC DAn displayed specific THD pathological features that matched patients’ phenotypes, such as lower levels of DA metabolites and reduced TH expression. We also observed and altered expression levels of DA genes, when compared to controls. Moreover, THD iPSC DAn showed abnormal morphology, including reduced neurite length and either an abnormal TH proximodistal gradient (THDA), or neurite arborization defects (THDB). Finally, treatment of THD iPSC DAn with L-Dopa rescued the neuronal defects and disease phenotype in neurons from type A THD, but not in those from the type B patient. Interestingly, L-Dopa treatment at the stage of neuronal precursors could prevent the appearance of alterations in THDB-iPSC derived DAn.

## Materials and methods

### Description of THD patients

Control and THD-specific iPSC lines were generated by the P-CMR[C] Stem Cell Bank. The subjects who participated in the study were patients attending the Neurometabolic Unit of Sant Joan de Déu Hospital (Barcelona, Spain) and the Neuropediatrics Department at the University Hospital Virgen de la Arrixaca (Murcia, Spain). One of the two THD patients has a Type A phenotype (homozygous for TH p.Arg233His mutation) whereas the other one has a Type B phenotype (heterozygous compound for TH p.Arg328Trp and p.Thr399Met mutations). Detailed clinical information of both patients is provided in **Table 1.**

**Table 1:**
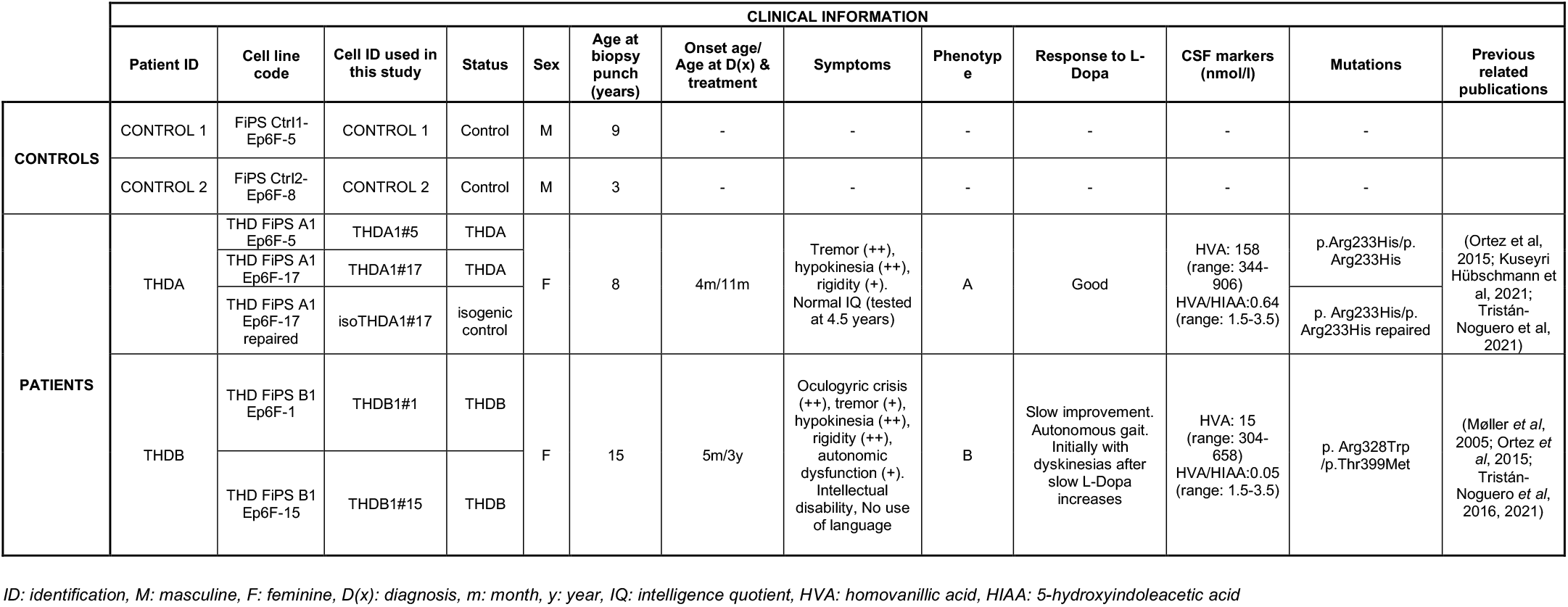
iPSC lines information of the clones used in the study and demographics of unaffected individuals and THD patients

### Generation and characterization of iPSC lines

Dermal fibroblasts were reprogrammed to iPSCs using non-integrating episomal vectors expressing Oct4, Sox2, Klf4, c-myc, p53shRNA and Lin28, as previously described (Kuebler *et al,* 2017) to generate 2-4 independent iPSCs clones per individual. Lines of control and patient-specific iPS cells were maintained in mTeSR™ medium (STEMCELL technologies) and passaged once a week with EDTA onto Matrigel-coated (Cultek) plates (Life technologies). Absence of episomal expression and endogenous pluripotency markers were assessed as previously reported (Kuebler *et al,* 2017). In vitro differentiation towards endoderm, mesoderm and neuroectoderm was performed essentially as previously described (Martí *et al,* 2013). The iPSC lines generated and characterized in this study have been registered and deposited in the Spanish Stem Cell Bank. iPSC lines information is provided in **Table 1**.

### Generation of CRISPR/Cas9 and gene edition in iPSCs

To repair the R233H mutation, mutant iPSCs were gene edited using CRISPR/Cas9. Three CRISPR/Cas9 gRNAs targeting the p.Arg233His mutation site in the TH gene were designed together with a ssODN (SIGMA) carrying the wild type allele and two additional synonymous mutations to eliminate the PAM sequence and to introduce a de novo HindIII to help with the molecular screening (GGGTCCCGAGCGCAGGGGCCCCTCACTGCCTGTACTGGAAGGCGATCTCAGCAATAAGCTTTCTGCGCTGGCGGTACACCTGGTCCGAGAAGCCCTGAGGG). The T7 endonuclease I assay (New england biolabs) was performed to assess which gRNA had the highest cleavage efficiency. The best performing gRNA (TH_ex6_gRNA1: ACCAGGTGTACCGCCAGCAC) was kindly provided by Synthego.

The selected iPSC line (THDA1#17) was seeded at a density of 300,000 cells per well of a six-well plate. The next day, iPSCs were nucleofected with a preassembled gRNA and Cas9 protein RNP complex (ThermoFisher) using the 4D AMAXA nucleofector (Lonza). Single colonies were screened by PCR followed by HindIII digestion (Thermo scientific). Those clones with a positive HindIII digestion were sequenced by Sanger sequencing. Clones presenting bi-allelic recombination and the desired genotype were expanded, cryopreserved and karyotyped. The maintenance of the expression of the pluripotency marker and the ability to obtain the three germ layers was verified by immunofluorescence.

### iPSC differentiation to DA neurons

Directed differentiation of iPSC onto DAn was carried out following a 30-days protocol based on DAn patterning factors and co-culture with mouse PA6 feeding cells to provide trophic factor support (Sánchez-Danés *et al,* 2012). Briefly, iPSCs were mechanically aggregated to form embryoid bodies (EBs). EBs were cultured 10 days in suspension in N2B27 medium, consisting of DMEM/F12 medium (GIBCO), Neurobasal medium (GIBCO), 0.5x B27 supplement (GIBCO), 0.5x N2 supplement (GIBCO), 2mM Ultraglutamine (Lonza) and penicillin-streptomycin (Lonza). In this stage, N2B27 was supplemented with SHH (100 ng/ml, Peprotech), FGF-8 (100 ng/ml, Peprotech), and bFGF (10 ng/ml; Peprotech). Neural progenitor cells (NPCs) were then seeded on top of PA6 for 21 days in N2B27 medium, as described (Sánchez-Danés *et al*, 2012). For L-Dopa and carbidopa studies, NPCs were seeded on the top of mouse PA6 cells and maintained in N2B27 medium, as described above. When cultures reached 20 days, treatment was initiated using L-Dopa (50 μM; 3,4-dihydroxy-L-phenylalanine, Sigma) and Carbidopa (12,5 μM; Sigma) in a ratio of 4:1 (Burbulla *et al*, 2017). Each compound was added when medium was changed, thus corresponding three times a week. DAn differentiations under treatment were kept in culture until they reached 30 days of culture, and a total of 10 days of treatment. For the early L-Dopa studies, the treatment was initiated at day 6 during EB’s neural induction and maintained until the end of differentiation (day 30) for a total of 24 days.

### iPSC differentiation to neural cultures not-enriched-in-DAn

To promote neuronal differentiation (not enriched in DAn) an existing protocol (Chambers *et al*, 2009) was adapted. Briefly, the iPSCs were maintained in mTeSR™ medium until confluence. EBs were generated and maintained in suspension for 48h. Next, the medium was changed to proneural (PN) medium and maintained for five days. PN medium consists of DMEM/F12 medium (GIBCO), Neurobasal medium (GIBCO), 0.5x B27 minus Vitamin A supplement (GIBCO), 0.5x N2 supplement (GIBCO), 2mM Ultraglutamine (Lonza), β-mercaptoethanol (Life Technologies) and penicillin-streptomycin (Lonza).

To obtain neural rosettes the EBs were seeded on poly-l-ornithine/laminin (POLAM) coated plates with PN medium supplemented with Noggin (200ng/ml; Peprotech) and SB431542 (10 μM; Tocris). After 8-12 days, neural rosettes were picked and expanded in new POLAM coated dishes. Neural rosettes were enzymatically dissociated and seeded in new POLAM-coated plates with PN medium supplemented with FGF2 (10 ng/ml; Peprotech) and EGF (10 ng/ml; Peprotech) to promote neural progenitor cells (NPC) generation and proliferation. After they had formed a homogenous cell population, the NPCs were further differentiated to GABA and glutamatergic neurons on POLAM-coated plates with PN medium during three to five weeks.

### Protein extraction and Western blotting

For pellet collection, cells were washed with cold PBS and mechanically detached from culture dishes and centrifuged for 5 min at 2000 rpm. Cell pellets were then resuspended in RIPA protein extraction buffer (Sigma) supplemented with protease inhibitor (Roche) on ice. Resuspended pellets were then sonicated in one pulse of 10 s at 10% amplitude on Branson Digital Sonifier® ultrasonic cell disruptor (Branson Ultrasonics Corporation) and let in a rotator during 1 hour at 4°C to homogenize proteins. The resulting supernatant was normalized for protein using Bradford method (Biorad). Total protein extracts were denatured in loading buffer for 5 min at 95°C. Each 15 μg sample of protein was separated using 8% SDS polyacrylamide gel electrophoresis (SDS–PAGE) and transferred to a nitrocellulose membrane (Sigma). The membrane was blocked with 5% not-fat milk in 0.1M Tris-buffered saline (pH 7.4)/Tween 0,1% (TBSTween) during one hour at room temperature and incubated overnight in TBS-Tween containing rabbit anti-TH (1:1000; Santa Cruz) at 4°C. After incubation with peroxidase-tagged secondary antibodies (1:2000; Amersham) for 1 hour at room temperature, membranes were revealed with ECL-plus chemiluminescence western blot kit (Amershan-Pharmacia Biotech). The intensity of the protein band was quantified using Fiji® software. The optical density value of each band was corrected by the value of beta-actin (1:2000; Proteintech).

### Intracellular dopamine ELISA

Cells were harvested with cold PBS and centrifuged for 5 min at 1200 rpm. The obtained pellet was diluted in 135 μl of sodium metabisulfite and EDTA both from Sigma, to prevent catecholamine degradation. Next, the pellet was sonicated at 10% amplitude for 60 s (10 s pulse alternating with 10 s of repose in ice) repeated three times. 115 μl of the 135 μl were kept for ELISA studies and the remaining volume was used to determine protein concentration with the Bradford method. Intracellular DA was measured using an ELISA kit for DA (LDN) in duplicates. This determination is a two-step process: first DA is extracted, it is acetylated and activated and then it is enzymatically converted and detected with a competitive ELISA assay. Corresponding DA levels were normalized to protein concentration determined previously.

### High Performance Liquid Chromatography (HPLC)

The supernatant was collected from neurons differentiated on the top of PA6 and kept directly at −80 °C until the moment of analysis. Before their analysis, medium samples were previously deproteinized with 50 μl of homogenization medium (100 ml miliQ H2O, 100 mg of sodium metabisulphite (Sigma), 10 mg of EDTA-Na (Sigma), 100 mg of cysteine (Sigma) and 3,5 ml of HClO4 concentrated (Scharlau, 70%)); centrifuged at 15000 rpm during 30 min at 4°C and the supernatant was filtered (0,45 μm, Millipore) for a posterior HPLC injection. The concentration of 3,4-Dihydroxy-L-phenylalanine (L-Dopa), dopamine (DA) and 3,4-dihydroxyphenylacetic acid (DOPAC) in supernatant (SN) samples was determined using an HPLC system consistent of a Waters 717 plus autosampler (Waters Cromatografia), a Waters 515 pump, a 5 μm particle size C18 column (100×46 mm, Kinetex EVO, Phenomenex) and a Waters 2465 amperometric detector set at an oxidation potential of 0.75 V. The mobile phase consisted of 0.15 M NaH2PO4.H2O, 0.57 mM 1-octane sulfonic acid, 0.5 mM EDTA (pH 2.8, adjusted with phosphoric acid), and 7,4% methanol and was pumped at 0.9 ml/min. The total sample analysis time was of 50 min and the L-Dopa, DA and DOPAC retention times were 2.06 min, 3.94 min and 4.25 min respectively. The detection limit was of 2-3 fmol (injection volume 60 μl). Corresponding DA metabolite content was normalized to protein concentration determined previously by the Bradford method detection.

### RT-qPCR analyses

Cells were harvested with cold PBS and centrifuged for 5 min at 2000 rpm. The pellet was kept directly at −80°C until the moment of analysis. Total mRNA was isolated using the RNAeasy Mini kit (Qiagen). 500 ng of total mRNA was used to synthesize cDNA with the SuperScript III First-Strand Synthesis System (Thermofisher). Quantitative PCR analyses were done in duplicate on 8 ng with PowerUp Syber Green Master Mix (Thermofisher) in an ABI Prism 7900HT Fast Real-Time PCR System (Applied Biosystems), using the following program: 50°C for 2 min, 95°C for 2 min, 95°C for 15 s, and 60°C for 1 min (two last steps repeated 40 cycles). All results were normalized to beta-actin and NSE (primers used in the study are listed in **Supplementary Table 1**).

### Immunocytochemistry

The iPSCs or iPSC-derived neurons were fixed with 4% paraformaldehyde solution for 20 min at RT. Antigen blocking and cell permeabilization were done using 3% donkey serum and 0.1% Triton X-100 in TBS for 2h at room temperature. Primary antibodies prepared in blocking solution were incubated for 48h at 4°C, while secondary antibodies (1:200; Alexa Fluor Series, Jackson Laboratories) were incubated for 2h at room temperature after washes. The primary antibodies used were mouse anti-ASMA (1:400; Sigma), goat anti-FOXA2 (1:50; R&D Systems), goat anti-Nanog (1:25; R&D Systems), mouse anti-OCT3/4 (1:25; Santa Cruz), rabbit anti-Sox2 (1:100; Thermofisher), rat anti-SSEA3 (1:10; Iowa), mouse anti-SSEA4 (1:10; Iowa), rabbit anti-TH (1:250; Santa Cruz), mouse anti-Tra-1-60 (1:100; Millipore), mouse anti-Tra-1-81 (1:100; Millipore) and mouse anti-Tuj1 (1:500; Covance). To visualize nuclei, samples were stained with DAPI (4,6-diamidino iamidino-2-phenylindole) and then mounted with PVA-DABCO.

### Neuronal quantification during differentiation

Immunostaining images were taken using a Carl Zeiss LSM880 confocal microscope and analyzed using Fiji® software to quantify the percentage of TH/DAPI, TUJ1/DAPI and TH/TUJ1 at day 30 (Schindelin *et al,* 2009). An average of five images were quantified for each ratio, and each differentiation was performed at least three times. Data points represent the average of at least three-independent experiments.

### Neurite, fiber density and TH proximodistal gradient quantification

The analysis was done at the end-point of the culture (30 days) on isolated iPSC derived DA neurons differentiated on top of PA6, either treated or not with L-Dopa and Carbidopa for 10 days and 24 days (early L-Dopa), fixed and stained for TH. Immunostaining images were analyzed with NeuronJ® software to quantify the number of neurites per neuron and neurite length for TH+ cells. Each neuron was analyzed in NeuronJ® and each trace was automatically measured and organized in order to obtain information for every single cell. An average of 5 images and ten neurons per image were analyzed for TH+ data for every iPSC-derived line. For the quantification of the fiber density, we generated a mask using ImageJ to delimit the area of the image occupied by TH + fibers which did not include nuclei (TH stained area). The area was corrected by the number of TH+ neurons present in each image (Prots *et al,* 2018; Ishikawa *et al,* 2016). To assess the TH proximodistal gradient, we only selected neurites longer than 60 μm, to rule out any confounding effect of the shorter total neurite length in THDA/THDB DAn, when compared to controls. Then the intensity of TH staining was measured at the distal end and compared with the one in the neuronal soma. Fluorescence intensity was calculated as previously described (Fitzpatrick, 2019) using the following formula:

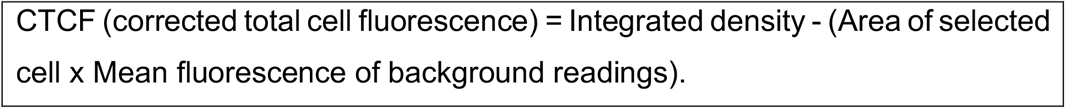

### Data Analysis and Statistics

For all experiments, one isogenic iPSC line, two control and two THD patientspecific iPSCs were used. Experiments were run at least in triplicate and averaged for each line. Normalized averages across multiple experiments were used to generate graphs of mean from each individual. We did not include in our analyses iPSC lines that did not grow well in each experiment.

Statistical n was performed solely on number of individuals per group, detailed in each figure and figure legend. The data were analyzed using analysis of variance (ANOVA), Kruskal-Wallis, or Student’s t/MannWhitney U tests according to the normality of comparisons and were plotted using Prism version 7.00 for Mac (GraphPad Software, La Jolla, CA, USA) with SEM error bars. ****p<0.0001, *** p< 0.001, **p<0.01 and * p<0.05. At least three independent experiments (n) were performed in all analysis.

This study includes no data deposited in external repositories.

## Results

### Generation of THD-specific iPSC lines

A total of seven iPSC lines from THD patients and healthy age-matched controls, along with a gene-edited counterpart, were used for the current study (see **Table1 and EV1**). Two clones of each cell line were thoroughly characterized and show a full reprogramming to pluripotency, as judged by colony morphology, alkaline phosphatase (AP) staining, expression of pluripotency markers (OCT4, NANOG, SOX2, SSEA3, SSEA4, TRA1-81 and TRA1-60), karyotype stability and silencing of episomal vectors (**Fig 1A-C**). The presence of TH mutations was confirmed by sequencing (**Fig 1D**). Finally, all iPSC lines differentiated into cell types of the three embryonic germ layers as indicated by expression of specific markers, namely FOXA2 (endoderm marker), αSMA (mesoderm marker) TUJ1 (ectoderm marker) (**Fig 1E**). Thus, iPSC generated from THD patients or from healthy individuals display *bona fide* pluripotent stem cell features, and were indistinguishable in all test performed, with the exception that THD-iPSC lines carried the TH mutations.

**Figure 1.**
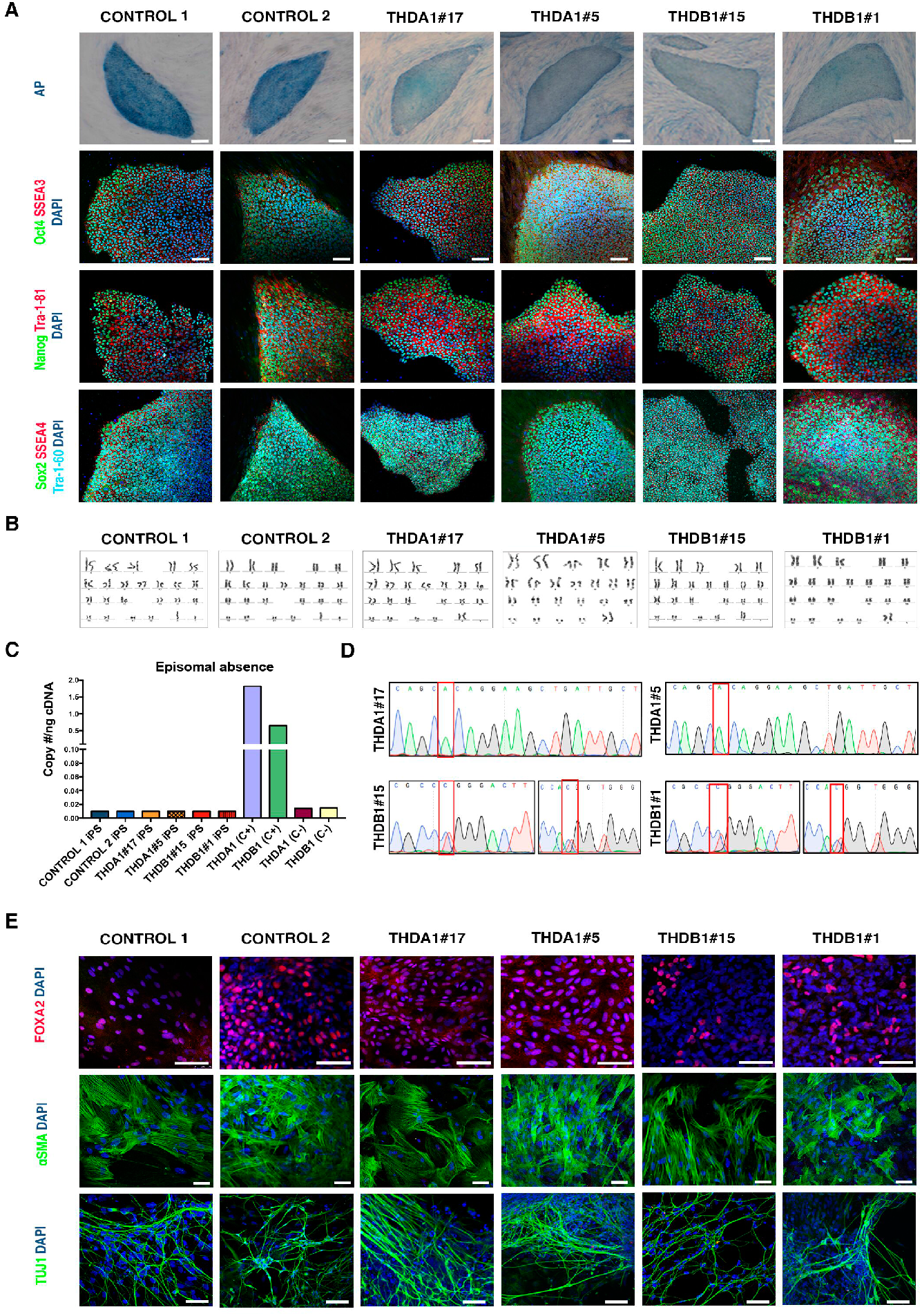
THD patient and healthy control iPSCs lines show bona fide pluripotent stem cell features. **A** CONTROL 1, CONTROL 2, THDA1#17, THDA1#5, THDB1#15 and THDB1#1 iPSC stained for alkaline phosphatase (AP) activity and for the pluripotency-associated markers OCT4, NANOG, SOX2 (green), SSEA3, TRA-1-81, SSEA4 (red) and TRA-1-60 (cyan). **B** No karyotypic abnormalities were observed in all iPSCs at passage 15. **C** Copy of gDNA (#/ng) of the episomal vector. Positive control (C+): THD iPSC line nucelofected with GFP, negative control (C-): fibroblasts of each THD patient. **D** Direct sequencing of genomic DNA from THDA and THDB iPSCs identifying the c.698G>A and c.982C>T and c.1196C>T mutations, respectively. **E** Immunofluorescence analyses of controls (CONTROL 1 and 2), THDA-iPSC (THDA1#17 and THDA1#5) and THDB-iPSC (THDB1#15 and THDB1#1) differentiated in vitro show the potential to generate cell derivatives of all three primary germ cell layers including endoderm (stained for FOXA2, red), mesoderm (stained for α SMA, green) and ectoderm (stained for TUJ1, green). In A and E nuclei are counterstained with DAPI, shown in blue. Scale bars, 100 μm.

Additionally, we generated a gene-repaired isogenic control from the THDA1#17 iPSC clone carrying the Arg233His mutation (hereafter referred to as isoTHDA1#17) using CRISPR/Cas9 (**Fig EV1A-C**). We focused on the mutant line THDA1#17 because it is homozygous for the most frequent mutation (Arg233His) found in THD patients that can cause a Type A or Type B phenotype depending on factors not yet fully understood (Willemsen *et al*, 2010; Fossbakk *et al,* 2014). After full characterization of isoTHDA1#17, we found that CRISPR/Cas9-editing did not affect isoTHDA1#17 pluripotency, differentiation potential or karyotype stability (**Fig EV1D, E**).

### Generation of THD-specific DA neurons

For the directed differentiation of iPSC towards DA neurons, we next subjected all the iPSC lines to a 30-days differentiation protocol using DA patterning factors and co-culture with the mouse PA6 stromal cell line (Sánchez-Danés *et al,* 2012) (**Fig S1A)**. As a technical control, we differentiated iPSCs from the same subjects into neural cultures not-enriched-in-DAn using a different protocol which allows to generate mainly glutamatergic and GABAergic neurons throughout neuroepithelial progenitor cells (NPCs) (Chambers *et al,* 2009) (**Fig S1B)**. Neuronal and DA neuron differentiation were evaluated at the end stage of differentiation by quantitative RT-PCR to examine the expression of dopaminergic-associated genes, such as *TH, AADC* (aromatic L-amino acid decarboxylase), *D1DR* and *D2DR* (dopamine receptors 1 and 2), *DAT* (DA transporter) and *VMAT2* (vesicular monoamine transporter type 2) (**Fig S1C**). Overall, the results of these analysis revealed an up-regulation of these genes in DAn derived from all iPSC lines applying the DA enriched protocol when compared to NPCs and neural cultures not-enriched-in-DAn. These data indicate that all iPSC lines generated DA neurons with *bona fide* dopaminergic molecular characteristics (Hwang *et al,* 2010). Moreover, we found that DAn generated from control and THD iPSC were mature as judged by immunofluorescence analysis against neuron-specific class III-b-tubulin (TUJ1) tyrosine hydroxylase (TH) (**Fig 2A**, **Fig EV2)**.

**Figure 2.**
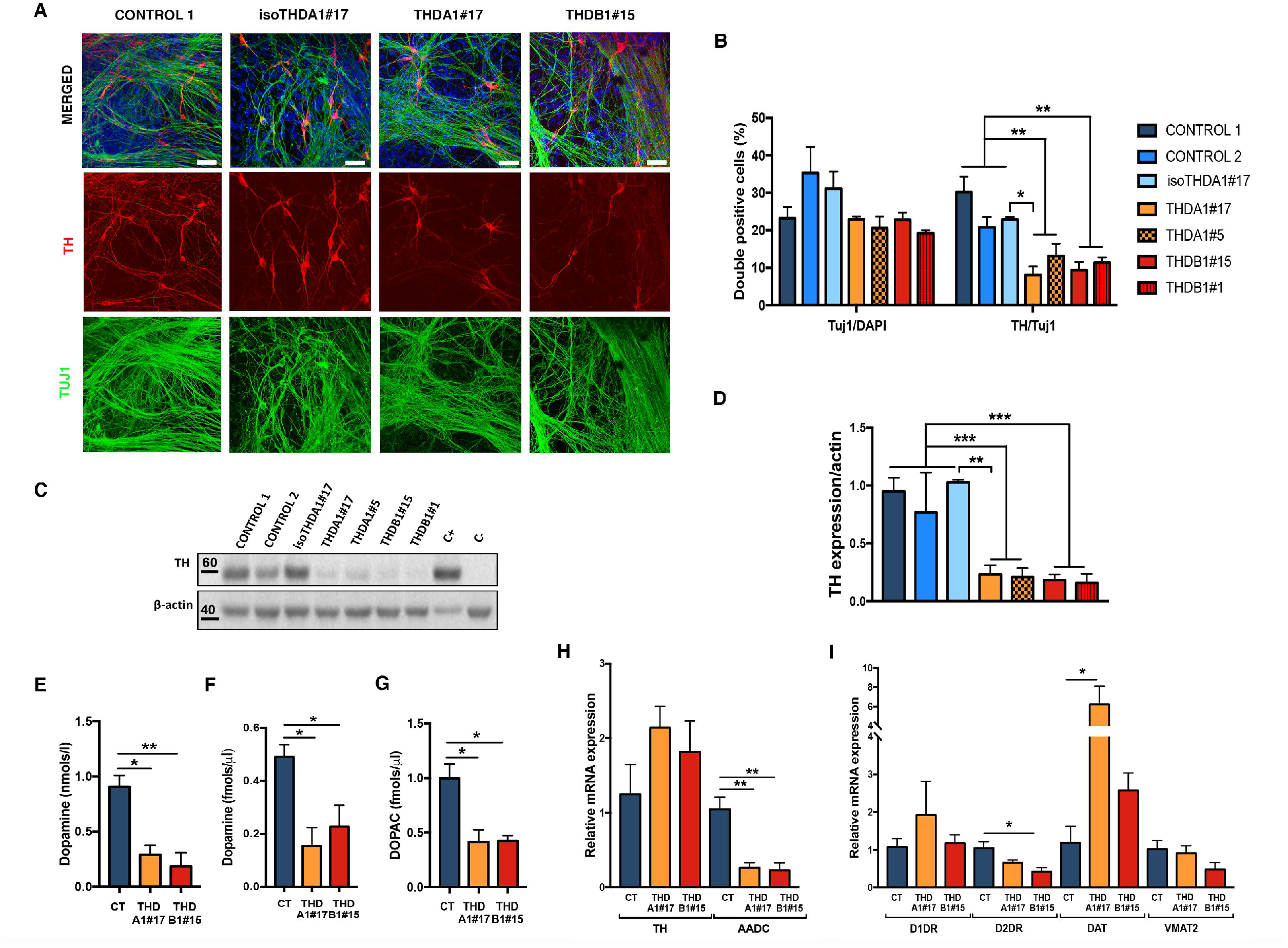
Dopaminergic neurons derived from THD-iPSCs recapitulate the disease phenotype. **A** Representative immunofluorescence (IF) images of CONTROL 1, isoTHDA1#17, THDA1#17 and THDB1#15 neuronal cultures (TH in red, Tuj1 in green, DAPI in blue) at day 30 of differentiation. **B** Quantification of Tuj1/DAPI and TH/Tuj1 ratios in all cell lines (CONTROL 1, CONTROL2, isoTHDA1#17, THDA1#17, THDA1#5, THDB1#15 and THDB1#1). **C** Western blot analysis for determining TH expression in each cell line; positive control (C+): mesencephalon, negative control (C-): fibroblasts. **D** Quantification of western blot results (normalized by beta-actin expression). **E** ELISA quantification of intracellular dopamine levels (nmols/l) in CONTROL 1, CONTROL 2, THDA1#17 and THDB1#15. **F** HPLC quantification of dopamine levels (fmols/μl). **G** HPLC quantification of DOPAC levels (fmols/μl). **H** Relative mRNA expression of dopaminergic enzymes (TH and AADC) in CONTROL 1, CONTROL 2, THDA1#17 and THDB1#15. **I** Relative mRNA expression of other DA genes expression (dopamine receptors, DAT and VMAT2). Scale bars, 20 μm. Number of experiments = 3 per line. Data are expressed as mean ± SEM. ANOVA or Kruskal-Wallis tests were performed for multiple comparisons. Unpaired twotailed Student’s t test or Mann-Whitney U test was used for pairwise comparisons. ***p<0.001 **p<0.01 *p<0.05.

We then investigated whether the presence of THD mutations impacts on TH expression levels by counting the number of TH-expressing cells in cultures enriched for dopaminergic neurons. After detailed analysis, we found less TH-immunoreactive cells (~5 to 10% p=, p<0.01) in THD neuronal cultures compared with control and isogenic iPSC-derived cells (~25 to 30% of all Tuj1+ cells). However, no differences were observed in the numbers of Tuj1+ neurons generated from all iPSC lines (**Fig 2A, B** and **Fig EV2**), suggesting that the enzymatic defect may specifically affect the production of DAn. To further confirm these results, we tested the expression levels of the TH protein and found significantly lower levels of TH expression in both THDA and THDB neuronal cultures compared to those obtained from control neuronal cultures (p< 0.001). **(Fig 2C, D**).

We next attempted to examine the functionality of the DAn differentiated from THD patient-specific iPSC, by determining the intracellular and released dopamine (DA) and its metabolite 3,4-dihydroxyphenylacetic acid (DOPAC) in cultured THD and control neurons. In all patient derived cultures, we found low intracellular dopamine levels compared to those measured in the control and isoTHD neuronal cultures (p<0.05 for THDA and p<0.01 for THDB) (**Fig 2E**). Moreover, analysis of DA release from media collected from THDA and THDB neuronal cultures, revealed low levels of extracellular dopamine and its metabolite 3,4-dihydroxyphenylacetic acid (DOPAC), compared to control and isoTHD cultures (p<0.05) (**Fig 2F, G**). These data confirm the severe effect of TH mutations in those neurons in which the reduced enzyme activity significantly affected both DA production and release. Interestingly, the use of the isoTHDA1#17 neuronal cultures restored the previous phenotype, confirming the dependence of the latter on the mutation (**Fig 2A-G**).

Finally, we investigated whether lower TH expression levels in THD dopaminergic cultures could impact on the expression of other DA-related genes. Overall, quantitative RT-PCR for dopamine-related gene expression revealed significantly lower *AADC* expression levels in both THD cultures (p<0.01; **Fig 2H**), lower *D2DR* expression levels in THDB cultures (p<0.05) and higher *DAT* expression levels in THDA neuronal cultures (p<0.05) (**Fig 2I**).

Taken together, these data reveal that our new THD-iPSC model recapitulates most phenotypes observed in patients allowing investigation of the THD pathogenesis in disease-brain cells.

### Morphological alterations of THD-iPSC-derived DA neurons

After establishing DAn-enriched cultures from both control and THD patient specific-iPSC, we evaluated whether there were any differences in neuronal phenotype between DAn derived from controls and those from patients’ iPSC lines. We noted that THD iPSC-derived DAn developed a range of abnormal morphologies over 30 days in culture that were not observed in control iPSC-derived DAn (**Fig 3A** and **EV3A**). Specifically, while DAn differentiated from control iPSC showed long neurites with complex dendritic arborization, DAn from THD-iPSC had fewer and simpler processes (**Fig 3B** and **EV3B**). To rule out any subjectivity in the attribution of the differences observed between these cultures, we directly measured the number and length of neurites. To this end, using high-power confocal images we randomly picked single TH-stained neurons from THD patient-specific and control iPSC lines (**Fig 3B** and **EV3B**; see also (Chu *et al,* 2009)). These analyses confirmed that the neurite length of control-iPSC-derived DAn was significantly higher than those of DAn differentiated from THDA and THDB iPSC (p<0.05) (**Fig 3B, C** and **EV3C**). Interestingly, the role of patients’ mutation was further confirmed in DAn differentiated from gene repaired iPSC, as the isoTHDA1#17 iPSC had longer total neurite length when compared to THDA1#17 (p<0.05) (**Fig 3B, C**).

**Figure 3.**
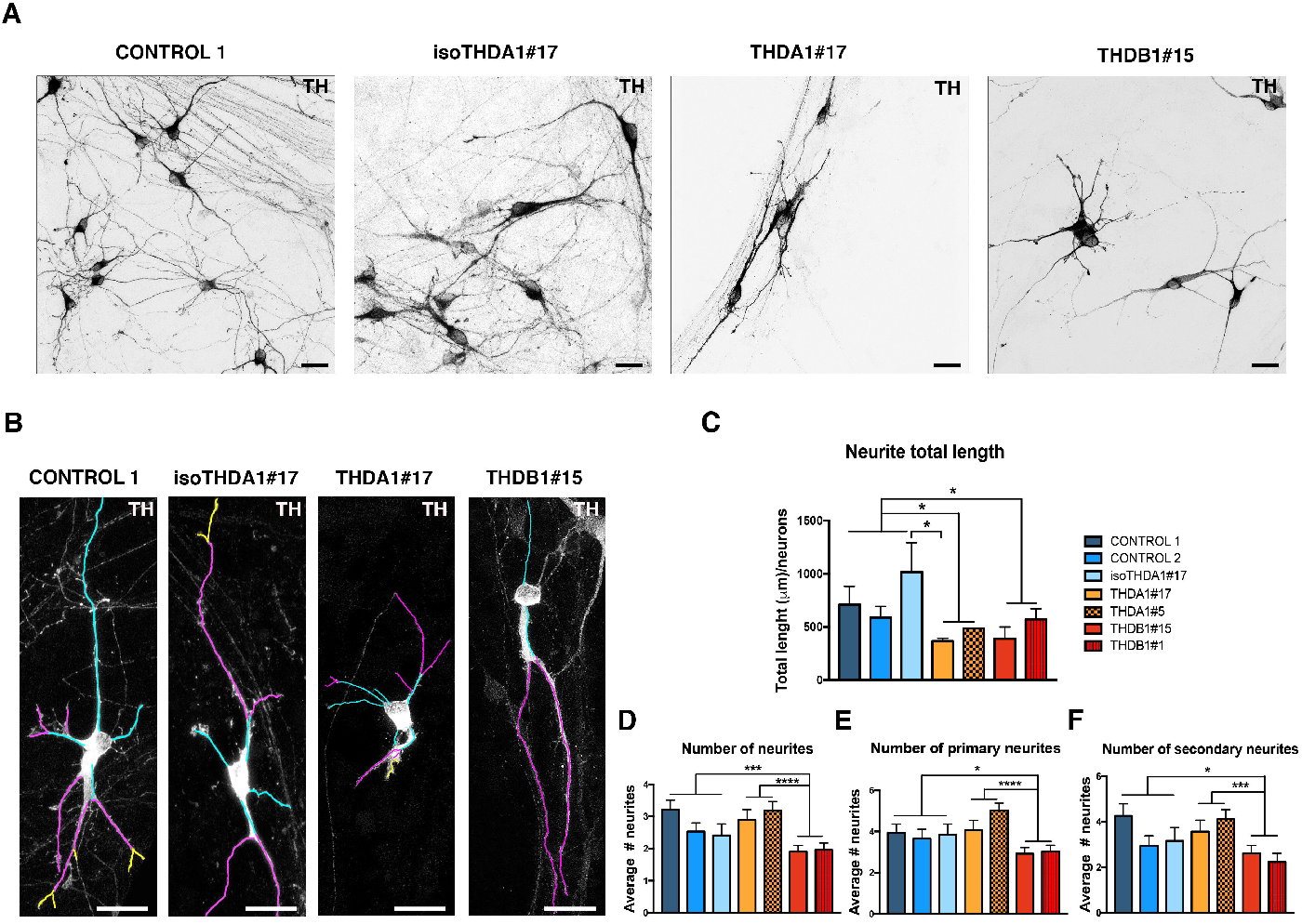
THD iPSC-derived TH+ neurons are morphologically abnormal. **A** Immunocytochemistry images of neuronal cultures (TH in black) of CONTROL 1, isoTHDA1#17, THDA1#17 and THDB1#15 cell lines. **B** Representative images of the tracing analysis (TH in white, primary neurites in blue, secondary in magenta and tertiary in yellow) in the same cell lines. **C** Quantification of the neurite total length in all cell lines (CONTROL 1, CONTROL2, isoTHDA1#17, THDA1#17, THDA1#5, THDB1#15 and THDB1#1). **D** Number of neurites. **E** Number of primary neurites. **F** Number of secondary neurites. Scale bars, 20 μm. Number of independent experiments = 3. Data are expressed as mean ± SEM. ANOVA or Kruskal-Wallis tests were performed for multiple comparisons. Unpaired two-tailed Student’s t test or Mann–Whitney U test was used for pairwise comparisons. ****p<0.0001 ***p<0.001 *p<0.05.

Additionally, THDB DAn showed fewer neurites when compared to controls and THDA DAn (p<0.001 and <0.0001, respectively) (**Fig 3B, D and EV3B**). Specifically, we observed fewer primary, secondary and tertiary neurites in DAn derived from THDB-iPSC compared to neurons derived from controls and THDA-iPSC (p<0.05 for primary and secondary neurites and p<0.01 for tertiary neurites when compared to control neurons; and p<0.0001 for primary neurites and p<0.001 for secondary neurites when compared to THDA neurons) (**Fig 3B, E,F and EV3B, C**).

Overall, our data suggest that although THD-specific iPSCs differentiated normally into DA neurons, they unveil morphological alterations upon 30 days in culture.

### THDA DA neurons show abnormal TH proximodistal gradient in neurites

Moreover, we observed that THDA DAn showed a distribution pattern of TH different from that of control and THDB neurons judged by TH immunofluorescence with anti-TH antibody (**Fig 3A, Fig EV3A and Fig 4A**). We validated this observation in subsequent, more detailed analyses for which we only selected neurites longer than 60 μm, to rule out any confounding effect of the shorter total neurite length in THDA and THDB DAn, when compared to controls. In the control and THDB DAn, the intensity of TH staining gradually decreased along the proximodistal axis of neurites, whereas this gradient was more accentuated in neurites of THDA DAn. We calculated the ratio between TH intensity values at the distal end of neurites and at the neuronal soma. TH intensity ratios in THDA DAn were statistically significantly lower than those of control and THDB DAn (**Fig 4A, B**), suggesting that THDA DAn fail to transport TH to the neurite end. This finding is consistent with the reduced stability of Arg233His mutant TH enzyme (Fossbakk *et al,* 2014), as well as with the decline in TH transport along the nigrostriatal axons described in the THD mouse model (Korner *et al*, 2015). Moreover, the abnormal TH distribution in THDA DAn neurites was normalized in isoTHDA1#17 (**Fig 4A, B**), implicating the role of TH Arg233His mutation in this disease-related phenotype.

**Figure 4.**
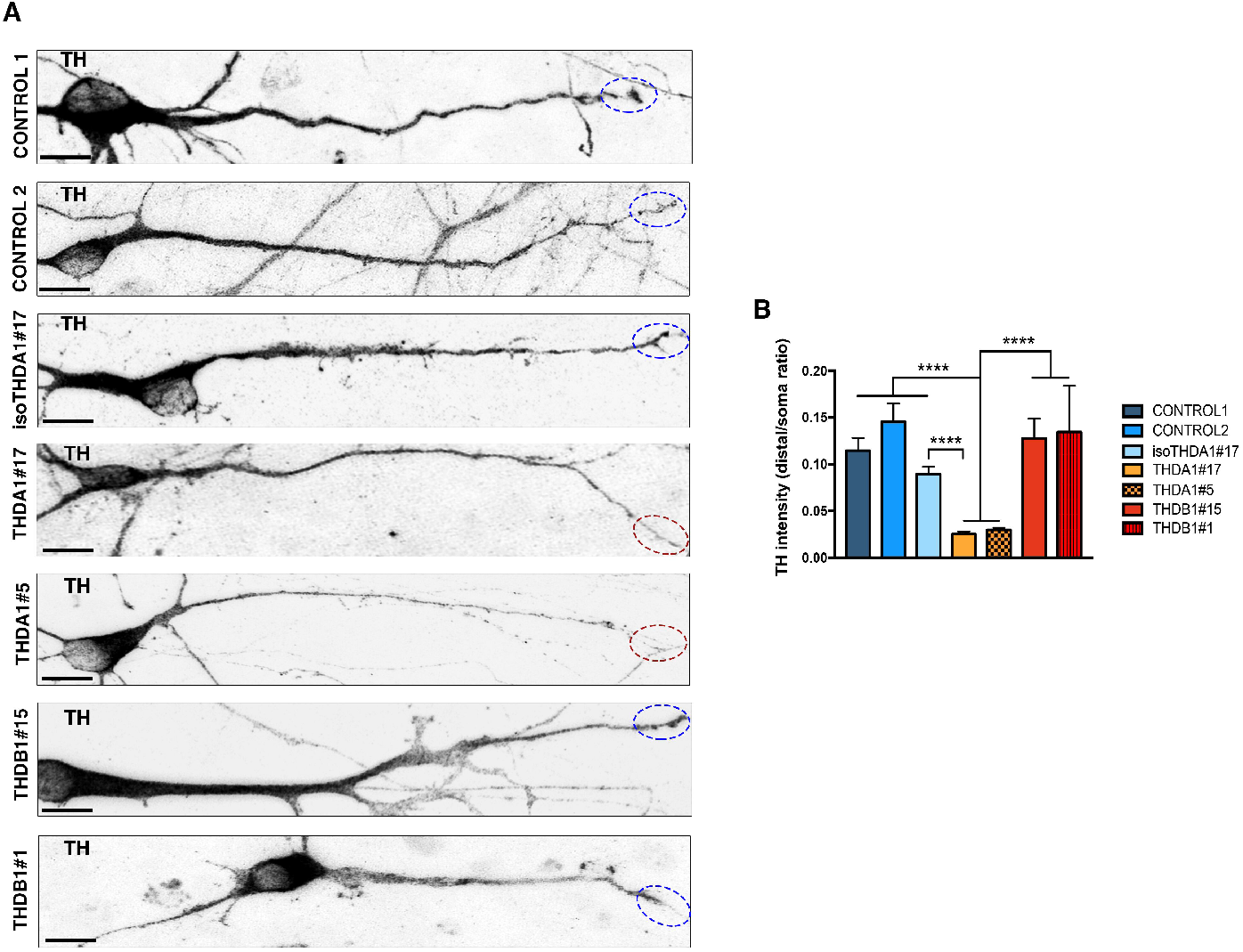
TH abnormal proximodistal gradient in THDA DAn neurites. **A** Representative images of DA neurons from CONTROL 1, CONTROL 2, isoTHDA1#17, THDA1#17, THDA1#5, THDB1#15 and THDB1#1 iPSC, stained for TH (black), circled neurite end. **B** Ratio quantification of TH intensity on the distal end of the neurite and the neuronal soma, in all cell lines (CONTROL 1, CONTROL2, isoTHDA1#17, THDA1#17, THDA1#5, THDB1#15 and THDB1#1). Scale bars, 10 μm. Number of independent experiments = 3. Data are expressed as mean ± SEM. Kruskal-Wallis test was performed for multiple comparisons. Unpaired two-tailed Student’s t test or Mann-Whitney U test was used for pairwise comparisons. ****p<0.0001.

### L-Dopa and Carbidopa treatment rescue the THD-specific phenotypes in THDA iPSCs-derived DA neurons

We next sought to obtain proof-of principle that L-Dopa treatment of THD patient-specific iPSC-derived DAn could rescue the morphological changes described above. For this purpose, we used L-Dopa together with Carbidopa (hereafter referred to as L-Dopa), because that is the way it is administered to patients in order to inhibit the AADC enzyme and avoid the peripheral L-Dopa metabolism (Willemsen *et al,* 2010; Furukawa & Kish, 2017). At the end of the 30-day differentiation protocol, that coincide with the end point treatment (Burbulla *et al*, 2017), the number of DAn was assessed by immunofluorescence for expression of TH and unbiased counting (**Fig 5A**). Treated THDA neuronal cultures showed significant more TH + neurons as judged by the TH/Tuj1 ratio (p<0.05) (**Fig 5B, C**). Fiber density was also increased in treated DAn derived from THDA iPSC (p<0.01) (**Fig 5B, D**). Moreover, we observed an increase in TH protein expression levels in treated THDA neuronal cultures (p<0.05) (**Fig 5E, F**). However, none of the previously described THD phenotypes were rescued in treated THDB neuronal cultures (**Fig 5B-F**).

**Figure 5.**
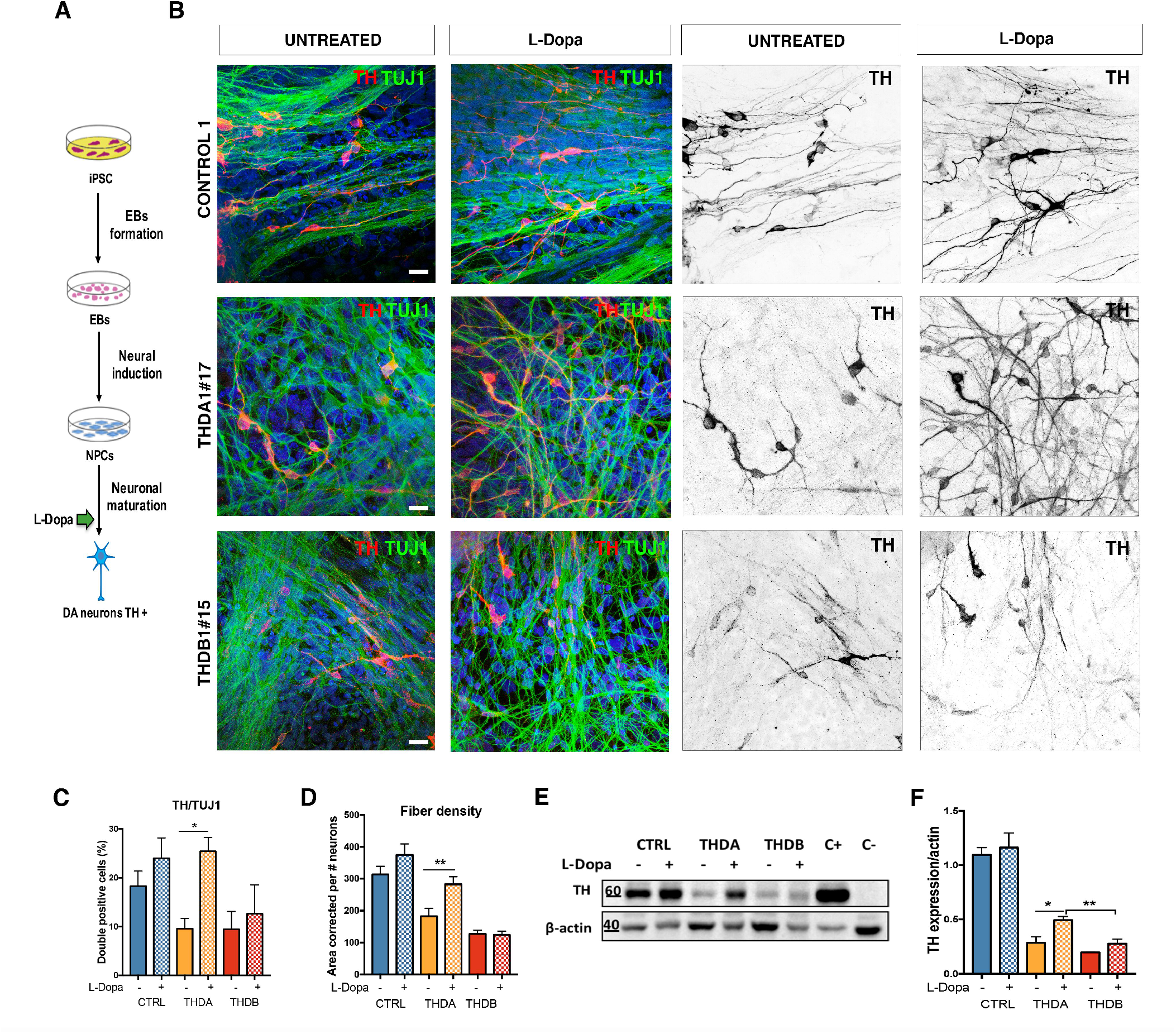
L-Dopa and Carbidopa treatment rescues the described phenotype only in THDA. **A** Schematic representation of treatment administration. **B** Immunocytochemistry images of CONTROL 1, THDA1#17 and THDB1#15 neuronal cultures (TH in red, Tuj1 in green and DAPI in blue) and in black and white to show TH staining. **C** Quantification of the TH/Tuj1 ratio. **D** Quantification of the fiber density. **E** Western blot of TH expression after L-Dopa treatment; positive control (C+): mesencephalon, negative control (C-): fibroblasts in CONTROL 1, THDA1#17 and THDB1#15. **F** Quantification of western blot results (normalized by beta-actin expression). Scale bars, 20 μm. Number of independent experiments = 3. Data are expressed as mean ± SEM. Unpaired two-tailed Student’s t test or Mann–Whitney U test was used for pairwise comparisons. **p<0.01 *p<0.05.

Neurite tracing analysis of treated control and THDA DAn revealed longer neurites (p<0.05 and <0.0001, respectively). Conversely, L-Dopa treatment did not rescue the abnormal arborization observed in THDB-iPSC derived DAn (**Fig EV4A, B**).

Next, we tested whether L-Dopa treatment could also increase DA metabolites levels in the treated neuronal cultures and found an increased intracellular DA levels in control and THDA neuronal cultures (p<0.0001 and <0.05, respectively) (**Fig EV5A**). Moreover, we found higher level of extracellular DA (p<0.0001 and <0.05, respectively) and DOPAC (p<0.01 and <0.001, respectively) in both control and THDA neuronal cultures (**Fig EV5B, C**). In contrast, none of the previously mentioned DA metabolites were detected at higher level in treated THDB DAn. Finally, as expected, we observed more L-Dopa in the supernatant media collected from control (p<0.01), THDA (p<0.05) and THDB (p<0.01) neuronal cultures (**Fig EV5D**). At the mRNA level, we found an increase in *AADC* mRNA expression levels in L-Dopa-treated control neuronal cultures (p<0.05) (**Fig EV5E**), whereas no statistical differences were observed in the expression of DA genes, such as *D1DR, D2DR, DAT* and *VMAT2,* in L-Dopa-treated control and THD neuronal cultures (**Fig EV5F**).

### Early L-Dopa treatment prevents THDB iPSC-derived DAn abnormalities

As DA is one of the earliest neurotransmitters being expressed during brain development, alterations in its signaling could affect the differentiation of specific subpopulations of neurons (Money & Stanwood, 2013). To directly assess the hypothesis that crucial events early in brain development might be responsible for the lack of treatment response in Type B (Willemsen *et al*, 2010), we established neuronal cultures from THDB-iPSC and assessed the effect of an early L-Dopa treatment. Specifically, L-Dopa and Carbidopa were added to the cultures starting from day six, at the stage of neuronal precursors, and kept until day 30 (**Fig 6A**), the time when THD phenotypes in THDB DAn were observed. Of note, the early L-Dopa treatment prevented the appearance of alterations in THD-B iPSC DAn, as showed by the increased number of TH-positive cells (p<0.05) (**Fig 6B, C**) and the restauration of the intracellular DA levels (p<0.05) (**Fig 6D**) as well as the extracellular levels of DA, DOPAC and L-Dopa (p<0.05) (**Fig 6E-G**). Moreover, the early L-Dopa treatment decreased the number of TH-positive cells with an abnormal morphology (**Fig 6H-K**). Importantly, these findings suggest the existence of a critical developmental window when new therapies could be successfully employed in THD.

**Figure 6.**
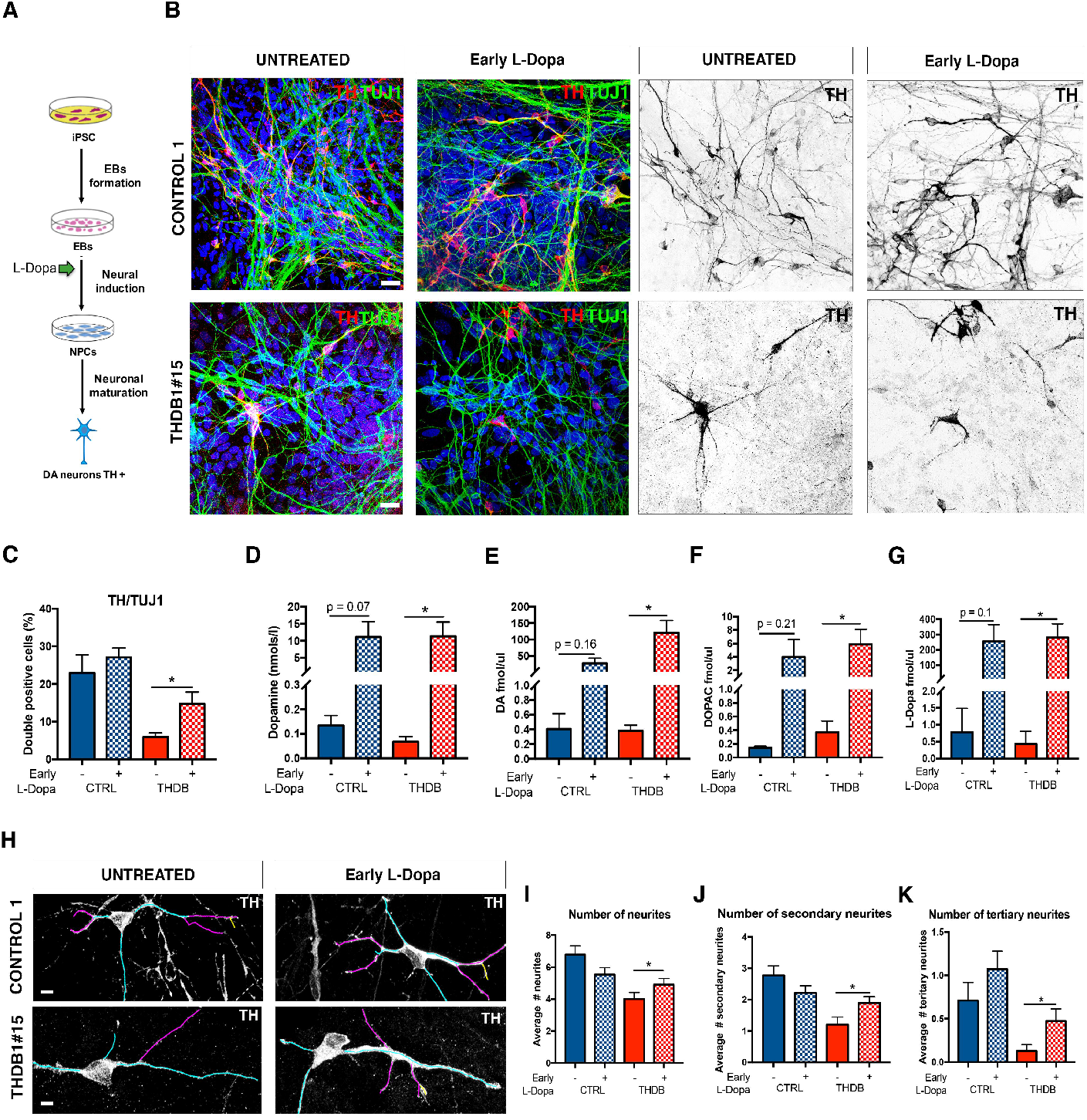
Early L-Dopa treatment prevents deficits in THDB. **A** Schematic representation of treatment administration. **B** Immunocytochemistry images of CONTROL 1 and THDB1#15 neuronal cultures (TH in red, Tuj1 in green and DAPI in blue) and in black and white to show TH staining. **C** Quantification of TH/Tuj1 ratio in the previous lines. **D** Elisa quantification of intracellular dopamine levels (nmols/l) in the CONTROL 1 and THDB1#15 lines before and after treatment. **E** HPLC quantification of dopamine levels (fmols/μl). **F** HPLC quantification of DOPAC levels (fmols/μl). **G** HPLC quantification of L-Dopa levels (fmols/μl) in CONTROL 1 and THDB1#15 lines. **H** Representative images of the tracing analysis (TH in white, primary neurites in blue, secondary in magenta and tertiary in yellow) of CONTROL 1 and THDB1#15 lines. **I** Number of neurites. **J** Number of secondary neurites. **K** Number of tertiary neurites. Scale bars, 20 μm. Number of independent experiments = 3. Data are expressed as mean ± SEM. Unpaired two-tailed Student’s t test or Mann–Whitney U test was used for pairwise comparisons. ****p<0.0001 **p<0.01 *p<0.05.

## Discussion

In this study, we describe the first iPSC model of THD with non-integrating episomal vectors of two THD patients, one with a mild phenotype (THDA) and another one with a severe phenotype (THDB). In addition, by CRISPR/Cas9 gene editing technique, we repaired the THDA iPSC line. Similar to previous studies modelling other dopamine neurotransmitter defects, we were able to obtain DAn from both control and THD patients independently of the disease status (Ishikawa *et al,* 2016; Rossignoli *et al,* 2021).

Here we show that our THD iPSC model recapitulates the disease phenotype associated with THD. As expected, THD patient-derived DAn showed a reduction in DA metabolites when compared to controls. The same was observed in mice models carrying mutations in the *TH* gene (Kobayashi *et al,* 1995; Althini *et al,* 2003; Tokuoka *et al,* 2011; Rose *et al,* 2015; Korner *et al,* 2015; Zhou *et al,* 1995). We also observed a reduced TH protein expression levels in both THDA and THDB derived DAn compared to controls. This effect was previously described in a *post-mortem* brain study of a Type B THD patient (Tristán-Noguero *et al,* 2016), in a PC12 cell line where *TH* mutations were exogenously expressed (Hole *et al,* 2015) and in THD mice models (Kobayashi *et al,* 1995; Korner *et al,* 2015). In this study we have also analysed different DA genes expression. We observed lower mRNA levels of *AADC* in both types of THD cultures and a reduction of *D2DR* mRNA expression only in THDB derived DAn but not in THDA. These findings are in line with previous literature: a *post-mortem* brain study of a Type B THD patient where reduced AADC and D2DR protein levels were observed (Tristán-Noguero *et al,* 2016). Moreover, a mouse model for Type A THD did not display lower expression of Dopamine receptors (Rose *et al,* 2015). It is generally agreed that in the presence of lower levels of DA, such as upon DAn denervation, up-regulation of dopamine receptors occurs (Turjanski *et al,* 1997). In our study, we have found lower levels of *D2DR* in THDB, but not in THDA. We hypothesize that low DA levels could regulate the expression of presynaptic dopaminergic receptors at an early developmental stage. Interestingly, the differences found between both receptors could cause an imbalance between direct and indirect pathways (Zeiss, 2005) and produce parkinsonian symptoms, which are usually worse in Type B patients (Iravani *et al,* 2012; Willemsen *et al,* 2010). Neurons derived from THDA-iPSC expressed higher mRNA levels of *DAT,* when compared to controls. This effect could be an adaptive behavior to use more efficiently the DA produced in *TH*-defective neurons by increasing DA uptake. Importantly this new human iPSC-based model generated from THD patients’ fibroblasts recapitulates some of the phenotypic aspects of human disease *in vitro.*

The most significant contribution of our work is the description of a new neuronal phenotype as we observed fewer and simpler processes in both THD-derived DAn. This was also observed in THD mice models, (Tokuoka *et al,* 2011; Korner *et al*, 2015; Rose *et al*, 2015) where lower TH signal in fibers, dendrites and projections was reported. We observed shorter neurites in both THD neurons which could be due to the lack of DA leading to a defect in dopaminergic neuron development (Li *et al,* 2015) and an abnormal arborisation in THDB neurons. DA is an important neurotrophic factor, which could boost a more complex neuronal morphology (Lieberman *et al*, 2018) to integrate synaptic information and promote neuronal plasticity (Arikkath, 2012). Most Type A patients show preserved cognitive and learning skills, which may be related to undamaged dendritic trees. In contrast, an important proportion of Type B patients have moderate to severe intellectual disability (Ortez *et al*, 2015; Willemsen *et al*, 2010). In this study, we found an abnormal neuronal dendritic tree in THDB DAn that could be explained by lower *D2DR* mRNA expression levels as this receptor has been described as a modulator of neuronal morphology in the frontal cortex and striatum (Money & Stanwood, 2013; Li *et al,* 2015; Lieberman *et al,* 2018). The abnormal TH proximodistal gradient in THDA DAn neurites would suggest a TH transport defect along the neuron that was already described in a Type B (TH p.Arg203His) mouse model (Korner *et al,* 2015). These morphological observations are important to elucidate the different Type A and B THD clinical phenotypes.

Importantly, CRISPR/Cas9 repair of the TH p.Arg233His point mutation abolished the THDA deficit, restoring both the number of TH + neurons and the TH protein levels to the ones observed in controls. An improvement in neuronal morphology was also observed. Finally, the abnormal TH proximodistal gradient was also normalized. Thus, the genome editing data confirms the robust consistency of mutation-related changes in THDA neuronal cultures.

It has been described that while Type A phenotype patients respond well to L-Dopa treatment (Willemsen *et al*, 2010), Type B phenotype patients respond poorly, have more severe symptoms (Iravani *et al,* 2012) and a major prevalence of L-Dopa-induced dyskinesias (Pons *et al,* 2013).

We tested the treatment with L-Dopa in neuronal cultures derived from one control and both THD patients. As expected, L-Dopa treatment fully rescued the neuronal phenotypes in THDA-derived neuronal cultures but not in THDB. The higher number of TH-immunoreactive cells observed in treated THDA cell culture has been also described in diverse animal models of Parkinon’s Disease (PD) where they show that either 6-OHDA or MPTP-lesioned animals treated with L-Dopa present more TH+ neurons. Thus, refusing the toxic effect of L-Dopa in the nigrostriatal population (Datla *et al*, 2001; Darmopil *et al*, 2008; DiCaudo *et al*, 2012). Higher TH protein expression levels could be due to a more stable TH enzyme due to its binding of L-Dopa or the catabolized DA. Related to this, previous *in vitro* experiments indicated that the specific *TH* mutation p.Arg233His could be affecting TH thermal stability (Fossbakk *et al,* 2014). The improved fiber density and neurite length only observed in THDA cultures may be explained by the effect of DA signaling through D2DR, whereas the lack of response of THDB neurons to L-Dopa could be due to lower basal levels of D2DR in THDB DAn. THDA neuronal cultures showed higher levels of DA metabolites after the L-Dopa treatment; surely the added L-Dopa is catabolized, increasing the concentration of DA and its metabolites. Thus, importantly, the THD iPSC model mimics not only the phenotype observed in THD patients but also the response to the treatment.

L-Dopa treatment was not able to rescue the deficits observed in THDB neurons, suggesting that crucial pathological events in THDB cells might be occurring early in neural development. To test the hypothesis, we administered L-Dopa at an early time point and test the response of THDB DAn. We observed that both the number and complex arborization of DAn increased. The first effect is already described in PD animal models treated with L-Dopa (Datla *et al*, 2001; Darmopil *et al*, 2008; DiCaudo *et al*, 2012) and the second one is observed in prefrontal cortex neurons with dopamine treatment (Li *et al*, 2015) and we hypothesize that the same could happen in the dopaminergic neuron population. The disease phenotype was also partially rescued with early L-Dopa treatment since DA metabolite levels were higher in THDB, indicating that the added L-Dopa is now metabolized, when provided during neuronal induction.

Overall, these results suggest that early treatment might prevent the alteration on DAn. However, translating it to the clinics would imply that this treatment should be given prenatally, which is not common since most diseases are not diagnosed until birth. This is the case of most inborn errors of metabolism, such as THD. However, an early treatment intervention has already shown promising results in guanosine triphosphate cyclohydrolase I (GTPCH, another enzyme involved in DA synthesis) deficiency (Brüggemann *et al,* 2012). Therefore, both the early diagnosis and early treatment of THD, may greatly improve patient outcome.

We should note that our results were obtained based on samples from two patients only; thus, more patients are needed to further validate our findings. Moreover, when working with stem cells, it should be kept in mind that variability between established lines may be found but in the present study all the recommended strategies to diminish the risk of cell variability were implemented (Escribá *et al*, 2021).

To sum up, our THD iPSC model is the first to recapitulate specific disease phenotypes and the response to treatment in both Type A and B phenotypes of THD. This model will help us better understand new molecular mechanisms of this disorder and it is a valuable platform to test different therapeutic approaches *in vitro* for the management of all THD patients.

## Acknowledgments

The authors are indebted to the THD patients who have participated in this study. The authors thank Bernd Kuebler and Begoña Aran from the P-CMR[C] Stem Cell Bank, for generating the THD iPSCs and who kindly provided the control iPSCs. We would like to thank Marcia Triunfol and João Mello-Vieira at Publicase for their help with this manuscript.

Research from the authors’ laboratories is supported by the European Research Council-ERC (2012-StG-311736-PD-HUMMODEL), the Spanish Ministry of Economy and Competitiveness-MINECO (RTI2018-095377-B-100 and PID2019-108792-GB-I00), Instituto de Salud Carlos III-ISCIII/FEDER (Red de Terapia Celular - TerCel RD16/0011/0024 and FIS PI15/01082 and P118/00111), Agencia Estatal de Investigación-AEI (Unidad de Excelencia María de Maeztu CEX2018-000792-M), AGAUR (2017-SGR-899), CERCA Programme/Generalitat de Catalunya and Fundació la Marató de TV3 (202012-31 and 202012-32). A.T.-N was partially supported by a PFIS pre-doctoral fellowship (FI14/00641) from the Instituto de Salud Carlos III-ISCIII and 2018BR-IRSJD-CdTorres from Sant Joan de Déu Hospital.

## Author roles

ATN, AGC and AC conceptualized and organized. ATN, IFC, CC, CB, LC were involved in the methodology. ATN, AGC and AC wrote the original draft. All authors were involved in writing, review and editing. LC, FA, RDJ, SI, MP, RA provided resources. ATN and CB performed statistical analysis. ATN, AGC and AC acquired funding. AR, AGC and AC supervised. All authors agreed to the authorship.

## THE PAPER EXPLAINED

### Problem

Tyrosine Hydroxylase deficiency (THD) is a neuropediatric disorder characterized by the lack of the tyrosine hydroxylase enzyme (TH) that is an important source of dopamine in the brain. The symptoms can resemble those of a movement disorder (phenotype called Type A), but can also include those of a severe, diffuse brain disorder (phenotype called Type B). Mild and moderate forms of THD show dramatic improvement when treated with L-Dopa, while patients with the severe form of THD do not respond to the treatment and present cognitive impairment. Here we have generated a new iPSC model for the better understanding of THD disease mechanisms and response to treatment.

### Results

We obtained induced pluripotent stem cells (iPSCs) from two THD patients: one with a mild phenotype (THDA) and another with the severe phenotype (THDB). Those iPSCs were differentiate into dopaminergic neurons (DAn), the cell type of interest of this disease.

Similar to what is observed in patients, THD model showed lower levels of dopamine and reduced TH expression when compared to controls. Moreover, THD DAn showed an abnormal neuronal morphology. This model also recapitulated the patient’s response to treatment as we were only able to rescue the defects in THDA but not in THDB. Interestingly, if the treatment was administered earlier in time, we could also prevent defects in THDB suggesting a precise time in brain development when treatment should be applied.

### Impact

This is the first iPSC-based model of THD that brings together THDA and THDB patients and recapitulates both disease features and response to treatment. Using this new model, we detected neuronal abnormalities that were never revealed before. We believe that this iPSC-derived model is a robust model to study THD disease mechanisms and will become an important tool to identify new therapies for the management of all THD patients.

### For more information

OMIM ENTRY: https://www.omim.org/entry/191290

iNTD (international working group of neurotransmitter disorders): http://intd-online.org/

DeNeu (Spanish patients association of neurotransmitters disorders): https://www.deneu.org/

Nord (organization of rare diseases patients). Rare disease database: https://rarediseases.org/rare-diseases/tyrosine-hydroxylase

## EXPANDED VIEW FIGURE LEGENDS

**Figure EV1. Isogenic control from p.Arg233His mutation carrier was generated and shows bona fide pluripotent stem cell features. A** A region of the exon 6 with the ssODN, the gRNA and the PAM sequence, the mutation repaired and the new restriction site for HindIII. **B** PCR of exon 6 (left) and digestion of this PCR product with HindIII (right) (green boxes show digested PCR products). **C** Sequence of a positive clone (3F) where the mutation was repaired, the sequence PAM is inactivated, and a new restriction site is introduced; all in homozygosis. **D** Representative images of isoTHDA1#17 iPSC stained positive for the pluripotency-associated markers OCT4, NANOG, Sox2 (green), SSEA3, TRA-1-81, SSEA4 (red) and Tra1-60 (cyan) and for the three primary germ cell layers including endoderm (stained for FOXA2, red), mesoderm (stained for αSMA, green) and ectoderm (stained for TUJ1, green). Nuclei are counterstained with DAPI, shown in blue. Scale bars, 100 μm. E Normal karyotype of isoTHDA1#17 iPSC at passage 25.

**Figure EV2. iPSC-derived neurons from THDA1#5, and THDB1#1 clones recapitulate the reduced number of TH+ neurons.** Representative immunofluorescence (IF) images of CONTROL 2, THDA1#5, and THDB1# iPSCs neuronal cultures (TH in red, Tuj1 in green and DAPI in blue) at day 30 of differentiation. Scale bars, 20 μm

**Figure EV3. iPSC-derived neurons from THDA1#5 and THDB1#1 clones show abnormal morphology. A** Immunocytochemistry images of neuronal cultures (TH in black) of the same lines. **B** Images of the tracing analysis (TH in white, primary neurites in blue, secondary in magenta and tertiary in yellow) in CONTROL 2, THDA1#5 and THDB1# neuronal cultures. **C** Number of TH tertiary neurites in all cell lines (CONTROL 1, CONTROL2, isoTHDA1#17, THDA1#17, THDA1#5, THDB1#15 and THDB1#1). Scale bars, 20 μm. Number of independent experiments = 3. Data are expressed as mean ± SEM. ANOVA test was used for multiple comparisons. **p<0.01.

**Figure EV4. L-Dopa and Carbidopa treatment rescue neurite total length in THDA. A** Representative images of the tracing analysis (TH in white, primary neurites in blue, secondary in magenta and tertiary in yellow) of CONTROL 1, THDA1#17 and THDB1#15 lines. **B** Quantification of the neurite total length of CONTROL 1, THDA1#17 and THDB1#15 untreated and treated cultures. Scale bars, 20 μm. Number of independent experiments = 3. Data are expressed as mean ± SEM. Unpaired two-tailed Student’s t test or Mann-Whitney U test was used for pairwise comparisons. ****p<0.0001 *p<0.05.

**Figure EV5. L-Dopa and Carbidopa treatment partially rescues THDA phenotype. A** ELISA quantification of intracellular dopamine levels (nmols/l) in CONTROL 1, THDA1#17 and THDB1#15 lines before and after treatment. **B** HPLC quantification of dopamine levels (fmols/μl). **C** HPLC quantification of DOPAC levels (fmols/μl). **D** HPLC quantification of L-Dopa levels (fmols/μl). **E** Relative mRNA expression (assessed by qRT-PCR) of DA enzymes (TH and AADC). **F** D1 and D2 dopamine receptors (D1DR and D2DR), DAT, and VMAT2 mRNA expression levels relative to Neural Specific Enolase (NSE) in CONTROL 1, THDA1#17, and THDB1#15. Number of independent experiments = 3. Data are expressed as mean ± SEM. Unpaired two-tailed Student’s t test or Mann-Whitney U test was used for pairwise comparisons. ****p<0.0001 ***p<0.001 **p< 0.01 *p<0.05.

